# Local goodness-of-fit measures for neural decoding

**DOI:** 10.64898/2026.11.10.765696

**Authors:** Sirui Zeng, Alison E. Comrie, Loren M. Frank, Uri T. Eden, Eric L. Denovellis

## Abstract

State-space models are widely used to relate neural activity to internal states that cannot be observed directly, but assessing whether a given model is appropriate for a dataset remains difficult. Current approaches, such as cross-validation, summarize model fit across an entire recording. They can reveal that a model is imperfect, but not *when* during the recording it fits poorly. We therefore lack tools for targeted model refinement. Here, we develop local goodness-of-fit measures that ask whether each spike is consistent with the model’s prediction based on all preceding spikes. Each spike, through the tuning of the neuron that fired, points to a range of likely states, just as the prediction does. A spike that narrows a broad prediction is consistent with it; one that points to states the prediction considers unlikely is not. Because the comparison needs only the prediction and the states each spike points to, it also applies to models that decode from unsorted spike waveforms. Using simulations in which model errors are known, we show that two consistency measures, based on the overlap of high-probability regions and on predictive checks, flag such inconsistent spikes, whereas the standard Kullback–Leibler divergence also flags spikes that merely narrow a broad prediction. We then apply the measures to a hippocampal recording from a rat performing a spatial foraging task. In these data, a model assuming continuous movement is expected to fail during immobility, when the hippocampal representation can jump to distant locations, so we compare it with a model that also allows such jumps. As expected, the overlap measure flags more spikes during immobility than during movement under the continuous model, and allowing jumps removes most of the continuous model’s flags. The consistency measures thus show *when* during a recording a state-space model fits poorly, providing tools for its targeted refinement.

## Introduction

Internal brain states related to memory, emotion, and attention play central roles in behavior. They arise from a time-varying interplay between the external world and internal processes and are not directly observable, making them difficult to characterize.

Recordings of neural activity, including spike trains, calcium signals, and voltage traces, provide experimental measurements influenced by these latent states. State-space models make it possible to relate these measurements to estimates of the latent states themselves. Specifically, these models provide a method for relating complex, high-dimensional neural activity to estimates of low-dimensional latent states and have provided insights into neural representations and dynamics associated with taste [1], spatial location [2], and aggression [3].

State-space models aggregate information about the latent states over time. They combine the information from the data at the current time (the normalized likelihood) with the accumulated information used to predict the latent state (the predictive distribution). This powerful combination enables robust estimation of the latent state, effectively balancing current information with the accumulated history of the system and prior knowledge.

Despite the wide use of state-space models in neuroscience, methods to validate them remain underdeveloped and underused. When validation is skipped or reduced to a single pass/fail verdict, a model that fails in one behavioral period but succeeds elsewhere is accepted or rejected wholesale, and the specific assumption responsible for a misfit is never identified—so the model is never refined in a targeted way. One reason models are not refined this way is that latent states are inherently difficult to evaluate [4]: in the absence of ground truth, model validity can only be inferred indirectly, for example by comparing relative likelihoods across models, the quality of reconstructed signals, or performance on downstream tasks.

Neural data are also dynamic, so model fit can vary substantially over time. Compounding this, the most common validation tools—cross-validation and “shuffle” permutation tests—typically summarize the entire recording in a single number: they report whether a model generalizes better than chance or better than a simpler alternative, but not *when* during a recording the model fits poorly or *which* of its components is responsible. Consequently, these methods can reject intentionally simplified yet scientifically meaningful models while offering little guidance for improving them. For example, analyses that decode rat movements from hippocampal place cells often ignore known features of the rat’s movement (such as momentum) and known features of neurons (such as refractoriness and bursting). These omissions can produce detectable model misfit even when the resulting decoded trajectories remain scientifically informative [5–7]. Assessing such models requires measures that indicate when a model fails and which of its assumptions is responsible.

State-space models already contain rich diagnostic information in the form of distributions used to estimate the state process at each time: the normalized likelihood, which reflects information from only the current neural observation, and the predictive distribution, which aggregates information from all past observations through the state-space model structure. One potential source of targeted, temporally local diagnostic information is a direct comparison of these two distributions. We do not expect them to be identical. The predictive distribution aggregates information from many past spikes, whereas the likelihood reflects only the current observation, so the two carry different amounts of information about the state and will typically differ in shape and spread.

What matters for model assessment is therefore not whether the two distributions differ, but whether they are *consistent*. We call them consistent when they concentrate their probability mass in overlapping regions of the state space, and inconsistent when those regions are largely disjoint. Consistency is distinct from accuracy: accuracy asks whether the decoded state matches an external ground truth, which is unavailable for a latent process, whereas consistency asks only whether the two internal sources of information about that state—the accumulated past and the current observation—concentrate in the same region. The two distributions may be broad or narrow, and the decoded state may lie far from the animal’s physical location, yet the distributions remain consistent as long as their high-probability regions overlap. When instead they provide inconsistent information, the mismatch can reveal where, when, and potentially how the model fails to capture the structure of the data. Examining these discrepancies therefore provides a direct, interpretable measure of model–data agreement—analogous to a local, time-resolved goodness-of-fit test.

To this end, we develop and validate a set of local goodness-of-fit measures, each built on the comparison between the predictive distribution and the likelihood. Because the most familiar way to compare two distributions is to measure their divergence, we include Kullback–Leibler (KL) divergence as a point of reference. Divergence measures, however, quantify how much two distributions differ rather than whether they are consistent, so they flag any mismatch, including one that arises from a broad predictive distribution or a narrow likelihood, even when the model captures the structure of the data well. We therefore develop two measures that target consistency: highest probability-density (HPD) overlap and a rank-based predictive *p*-value derived from a predictive check.

In simulated data with known misspecification, we show which misspecifications each measure detects and that the consistency measures are not triggered when the latent state departs from the animal’s physical position without misfit, as in replay. In a hippocampal recording, we then use them to locate where a decoding model fails and to guide its refinement. These results indicate that consistency is the right target for local goodness-of-fit assessment, and they suggest a path from binary model rejection (is this model good or bad?) toward an iterative, diagnostic approach to building interpretable theories of neural dynamics.

## Methods

### State-space models

The state-space paradigm is particularly well suited for neural data analysis because many neuro-science experiments focus on internal brain processes that are not directly observable. For example, hippocampal replay of spatial trajectories has been hypothesized to be related to mental exploration of space, but the trajectories being replayed are not directly observable by experimentalists because these trajectories are different from the spatial position of the animal [8]. In such cases, state-space models are used to simultaneously estimate the unobserved internal representations and their relation to neural activity patterns.

For a discrete set of time points, *t*_0_, …, *t_K_*, we define an unobserved state process *x_k_* with initial state *x*_0_, which influences a neural observation process *y_k_* at each subsequent time *t_k_*, *k* = 1, …, *K*. In the models analyzed here, *y_k_* comprises the spikes of each sorted unit in the *k*th time bin; in a clusterless model, each spike would instead be identified by its waveform features. We define a state-space model by combining a state-transition distribution *p*(*x_k_* | *x_k_*_−1_), which defines how the state evolves in time, with an observation likelihood *p*(*y_k_* | *x_k_*), which defines the relationship between the neural activity and the state. To simplify the problem, we assume that the state-transition distribution depends only on the most recent past value of the state and that the observation likelihood depends only on the current state value. Fig. 1a shows a schematic of this state-space model structure.

**Fig 1.**
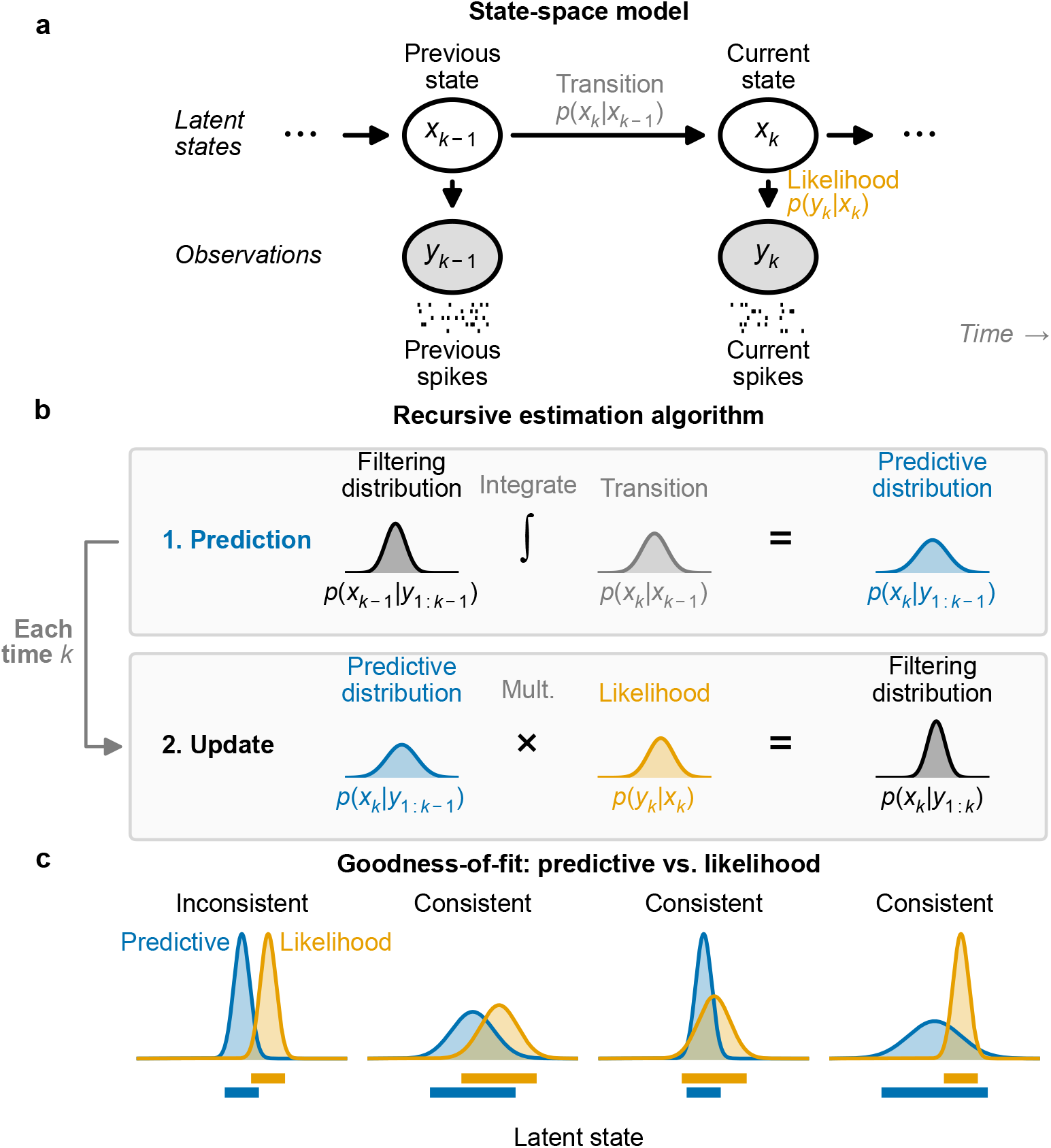
State-space model and local goodness-of-fit measures. **(a)** The graphical representation of the state-space model includes the state-transition model, *p* (*x_k_ | x_k−1_*), and the observation likelihood, *p* (*y_k_ | x_k_*). **(b)** These model components are sufficient to compute an iterative expression for the filtering distribution at time *t_k_*, *p (x_k_ | y*_1:_*_k_*), from the distribution at the previous time step. This involves first integrating the previous filtering distribution against the state-transition distribution to compute the one-step predictive distribution, *p* (*x_k_ | y*_1:_*_k−1_*), and then updating the predictive distribution with information from the most recent spiking by multiplying by the observation likelihood. **(c)** For each new spike observation, we compute the local goodness-of-fit by assessing the consistency of information between the predictive distribution and the normalized likelihood of that spike. Bars beneath each pair mark the high-probability region of each distribution. We consider the distributions inconsistent when they assign high probability to mostly disjoint subsets of the state space. When the regions with high probability overlap, the distributions are considered consistent, even if the distributions differ in their central tendency or certainty.

This state-space model structure allows for iterative estimation of the state process at each time step based on all the neural data up to that time (Fig. 1b). If the distribution of the state at time *t_k_*_−1_ has been computed, we can integrate it against the state-transition distribution to obtain a one-step predictive distribution, 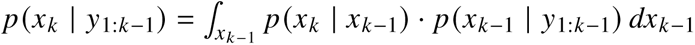 This predictive distribution combines the information from all previous observations to compute the distribution of the state at the next time step. We update this predictive distribution with the information from the most recent observation by multiplying by the observation likelihood to compute the filtering distribution, the posterior distribution of the state given all of the data up to the current time step: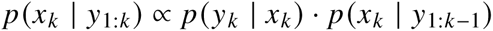

Common approaches for assessing the goodness-of-fit of state-space models include examining cross-validated error measures in the state process when the state is observable [9], cross-validated error measures in the observation process such as the mean-squared prediction error [10], and the distribution of prediction residuals [11]. Although these approaches can be useful, they may be difficult to interpret for neural data because intentionally simplified state-space models often exhibit some detectable lack of fit even when they remain scientifically useful.

Here, we take a different approach that relies on a set of local goodness-of-fit measures. Each measure compares the one-step predictive distribution with either the normalized likelihood of a single spike as a function of state or the spike itself. These measures therefore consider the consistency between information accumulated through the state-space model over past spiking and the information from the most recent spiking activity. Critically, we do not expect these distributions to align perfectly since they likely contain different amounts of information about the state process. Instead, we use the operational notion of consistency illustrated in Fig. 1c. We consider the distributions inconsistent when their probability masses are concentrated in mostly non-overlapping regions of the state space. We consider them consistent when their high-probability regions largely overlap, even if their central tendencies differ. We also regard the distributions as consistent when a broad high-probability region of one distribution contains a substantially narrower region of the other. This situation may arise frequently because the predictive distribution contains information accumulated from many previous spikes.

In the analyses here, “local” means that each measure is evaluated once per spike. The filter updates with the likelihood of everything observed in a time bin, including neurons that remained silent, whereas each measure compares the predictive distribution with the normalized likelihood of a single spike; bins without spikes therefore produce no diagnostic value. “Local” does not mean that the predictive distribution is restricted to recent data: it summarizes all observations through the preceding time bin via the recursive state-space filter. More generally, the same comparisons could use a filtering or smoothing distribution defined over another chosen window, including a window containing both past and future observations.

### Local goodness-of-fit measures

In this section, we introduce three measures of the local goodness-of-fit of the state-space model. All three are implemented in the open-source Python package `statespacecheck`, which can be applied to the output of other state-space decoders (see Discussion and Code availability).

All three measures are evaluated at the level of individual spike events. Let *C* be the number of recorded neurons, let *n_k_*_,*c*_ be the spike count of neuron *c* in time bin *k*, with **n***_k_* = (*n_k_*_,1_, …, *n_k_*_,*C*_), let 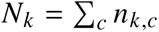 be the total population count, and let *y_k_*, *_j_* denote the mark of the *j*th event: the neuron identity for a spike-sorted model or the waveform features for a clusterless model. For a spike-sorted model, the observation *y_k_* is the count vector **n***_k_*, whose events carry the marks *y_k_*_,1_, …, *y_k_*_,*N*_*_k_*, so the observation likelihood *p*(*y_k_* | *x_k_*) is *p*(**n***_k_* | *x_k_*). For conditionally independent Poisson counts, it can be factored as

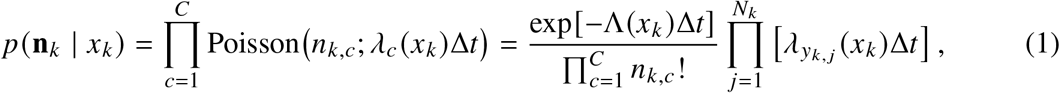

where Δ*t* is the bin width and *λ_c_* (*x_k_*) is the conditional firing rate of neuron *c* in the bin. Its expected count is *m_c_* (*x_k_*) = *λ_c_* (*x_k_*)Δ*t*. The total population rate is Λ(*x_c_*) = ∑*_c_ λ_c_* (*x_k_*), giving a total expected count of Λ(*x_k_*)Δ*t*. Equation (1) separates the exposure factor exp[−Λ(*x_k_*)Δ*t*], which is shared by the whole bin, from one event factor *λ_yk_*_, *j*_ (*x_k_*)Δ*t* for each observed event.

For the spike-sorted models analyzed here, HPD overlap and KL divergence compare the one-step predictive density

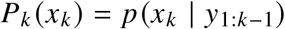

with the normalized single-event likelihood obtained from that event’s factor in Eq. (1):

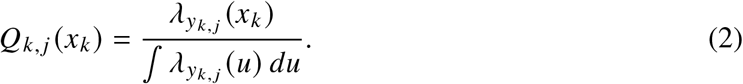

The common factor Δ*t* cancels in this normalization. *Q_k_*, *_j_* contains only the likelihood factor attributable to the selected event. It deliberately excludes the bin-level exposure term, silent neurons, and total spike count, which cannot be assigned to one particular event. Those aspects of fit require a complementary count or time-rescaling test. If several spikes occur in one bin, each event is evaluated separately against the same one-step predictive density *P_k_*. For a clusterless model, the analogous *Q_k_*, *_j_* is obtained by normalizing the marked event-intensity density over the state space.

#### Overlap between HPD regions

One way to assess the consistency between the predictive distribution and the observation likelihood is to examine the overlap between their highest probability-density (HPD) regions. The 95% HPD region of a distribution is the smallest set of states that together hold 95% of its probability: every state inside it is more probable than any state outside it, and for a distribution with several peaks it can consist of several separate pieces. We refer to the resulting measure as HPD overlap. A substantial intersection indicates that the distributions assign high density to some of the same state values. Minimal or no overlap indicates disagreement under this criterion. This measure provides an interpretable geometric summary of local consistency between the distributions.

We define the overlap between the HPD regions of the predictive distribution and the normalized single-event likelihood as

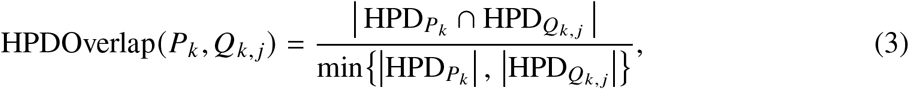

where HPD*_Pk_* and HPD*_Qk_*_, *j*_ are the highest probability-density regions of the predictive density and the normalized single-event likelihood, respectively, and | · | denotes the volume of the region. The numerator is the size of the overlap between these regions, and the denominator is the size of the smaller of the two HPD regions. This ratio is the Szymkiewicz–Simpson overlap coefficient applied to the two HPD regions. Unlike the Jaccard index, which divides by the size of the union and therefore penalizes any difference in region size, the overlap coefficient equals 1 whenever one region is nested inside the other, which is exactly the nesting-tolerant behavior our consistency criterion requires. This normalization ensures that the resulting value is in the range [0, 1], with 0 indicating no overlap and 1 indicating that one region is entirely contained within the other. For example, if the likelihood region is wider and therefore less informative than the predictive region but completely contains it, then the HPD overlap will be 1, suggesting that the two distributions are consistent under this criterion. Conversely, if the ratio is close to 0, it indicates little or no overlap, suggesting substantial differences between the ranges of state values inferred from the two distributions.

Fig. 2a illustrates the computation of the HPD overlap. We compute HPD regions for the predictive distribution (top) and the normalized single-event likelihood (middle), then divide the volume of their intersection by the volume of the smaller region (bottom).

**Fig 2.**
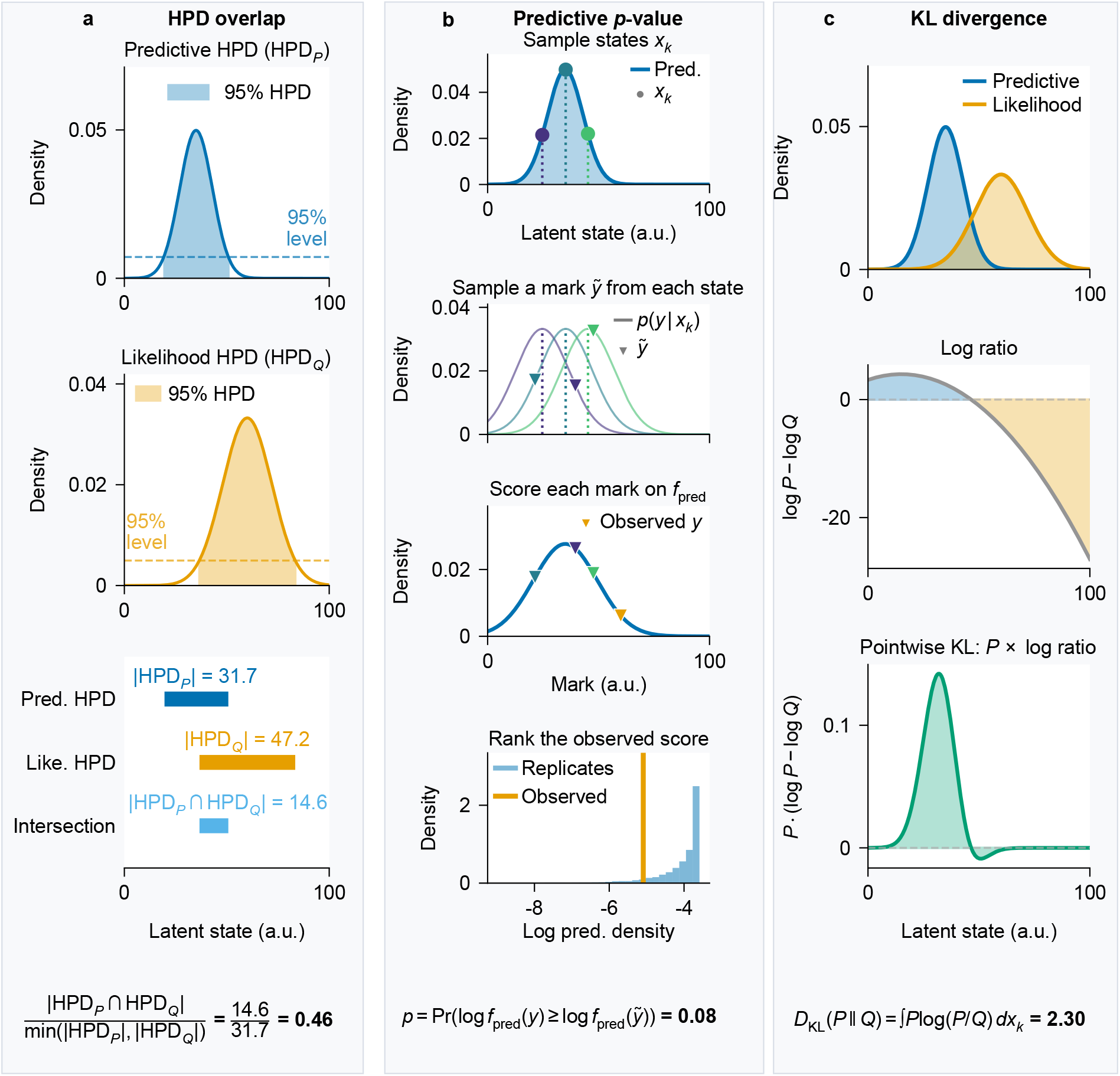
Computation of the three local goodness-of-fit measures. *P* is the one-step predictive distribution of the state and *Q* the normalized single-event likelihood of an observed mark *y* (a spike’s neuron identity or waveform features). **(a)** For HPD overlap, the volume of the intersection (bottom) of the 95% HPD regions of *P* (top) and *Q* (middle), bounded by the dashed levels, is divided by that of the smaller region. **(b)** The rank-based predictive *p*-value by sampling, for a continuous mark equal to the state plus noise. Illustrative states are drawn from the event-weighted predictive distribution (top). Each state generates a replicated mark from *p* (*y | x_k_*), the distribution of a spike’s mark given the state (second). Each mark is scored by its height on *f*_pred_, the average of those distributions over *P* (third), and *p* is the fraction of many replicates scoring no higher than the observed mark (bottom). For spike-sorted data, *p* is evaluated exactly. **(c)** For KL divergence, *P* (blue) and *Q* (orange) are compared (top) by the difference in log densities (middle), weighted by *P* (bottom) and integrated.

#### Predictive checks

The HPD overlap compares two distributions of the state: one based on the accumulated past information and the other based on the isolated information from the likelihood of the most recent spiking observation. Another approach is predictive checking, which compares the observation itself to a predictive distribution of that observation estimated from the model. This approach has been used before in other contexts [12–14] and has the advantage that it allows for the computation of *p*-values for individual observations, which are easy to interpret [15, 16].

Predictive checks for spiking data can be challenging to implement because, within sufficiently short intervals, a spike is typically less probable than no spike. One possible approach is to consider spike patterns over longer fixed intervals in which multiple spikes are expected. Here, we take a simpler approach of focusing only on spike times and measuring the probability that an observed spike comes from a particular neuron (for spike-sorted models) or has a particular set of waveform features (for clusterless models).

Using the event notation introduced above, the relevant observation distribution is the distribution of the spike’s mark for an event drawn from the model-implied marked point process. Because the *p*-value is evaluated once for each spike event, its predictive reference distribution corresponds to selecting an event rather than selecting a time bin that contains one or more events. Under the one-step predictive distribution *P_k_*, the expected number of events in the bin associated with a state region *dx_k_* is Λ(*x_k_*)Δ*t P_k_* (*x_k_*) *dx_k_*. The latent state associated with a randomly selected predicted event therefore has the event-weighted distribution

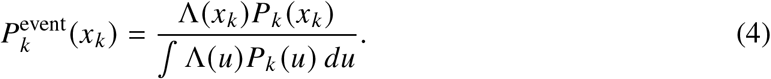

At states where Λ(*x_k_*) > 0, the probability that the event came from neuron *c* is

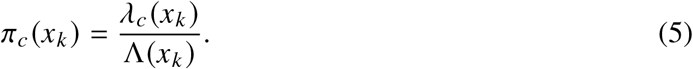

States where Λ(*x_k_*) = 0 receive zero mass under 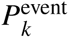 Combining the event-weighted state distribution with the conditional cell-identity distribution gives the predictive probability of cell identity:

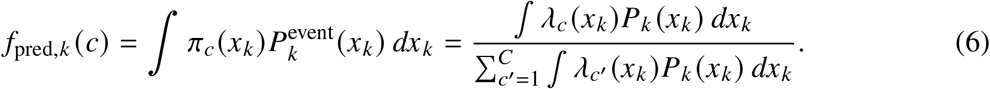

For a clusterless model, let *λ*(*y*, *x_k_*) be the event rate per unit mark volume at waveform mark *y*, so that Λ(*x_k_*) = ^∫^ *λ*(*w*, *x_k_*) *dw* is the total event rate and Λ(*x_k_*)Δ*t* is the total expected event count in the bin. The corresponding predictive mark density is

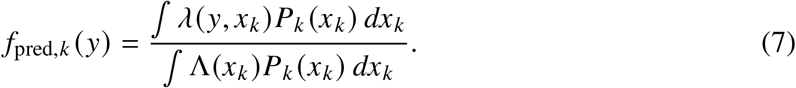

Equation (6) is the discrete spike-sorted form of Eq. (7). Let 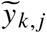 denote a replicated mark distributed according to *f*_pred,k_. We compute the probability that the observed spike is at least as likely as this replicate:

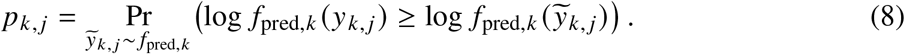

For the spike-sorted models considered here, 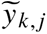 takes one of a finite set of values—the identities of the recorded neurons. We compute Eq. (8) exactly as

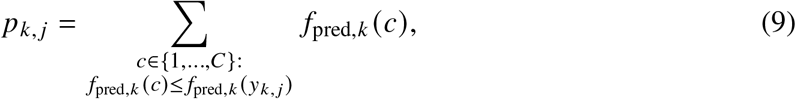

which sums the predictive probabilities of all cell identities at least as unlikely as the observed identity. Because *p_k_*, *_j_* ranks the observed spike against this predictive distribution of possible spikes, we refer to it as the *rank-based predictive p-value*.

Fig. 2b shows the same probability computed by sampling, the approach one would use for a continuous or otherwise intractable mark space. States are drawn from the event-weighted predictive distribution 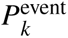 (top; in the schematic the total expected event count is constant across states, so it equals the one-step predictive distribution). Each state generates a replicated mark (second), each mark is scored by its predictive density *f*_pred,*k*_ (third), and *p* is the fraction of replicates scoring no higher than the observed mark (bottom). The spike-sorted results in this paper use the exact sum in Eq. (9). A *p*-value near zero indicates that the observed spike is unexpected given the past spiking and the model.

While HPD overlap explicitly assesses our definition of consistency in the state space, the predictive check probes a related notion in the observation space through the marginalization in Eq. (7). When the observation model is sufficiently informative about the state, we expect large *p*-values when the predictive distribution and likelihood match, but also when either distribution is narrower than the other, as long as they point to overlapping regions of the state space. This correspondence is not guaranteed: marginalization can map distinct state distributions to similar mark distributions, so a high predictive *p*-value does not by itself establish that the state distributions overlap. In the common case that the predictive distribution of the state is narrow relative to the likelihood, the predictive distribution of observations will be concentrated on the spikes that occur at the confidently predicted location, and the observed spike will have appreciable likelihood at that location. The observed spike is among those that the model anticipates and should therefore lead to a high *p*-value. In the less common case that the likelihood is narrow relative to the predictive distribution of the state, the predictive distribution of the observations will also be broad and will assign appreciable probability to a wide range of spikes. The observed spike is likely to fall within this anticipated range, so the *p*-value should also be high. A low *p*-value tends to arise when the observed spike is specifically unlikely relative to what the model anticipates, which corresponds to the likelihood concentrating in a region the accumulated past deems improbable.

#### KL divergence

Finally, one of the best-known measures of discrepancy between probability distributions is the Kullback–Leibler (KL) divergence [17], defined in this case as the expected excess logarithmic loss incurred when the normalized single-event likelihood is used to approximate the predictive distribution, with the expectation taken with respect to the predictive distribution:

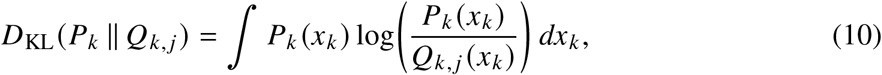

where *P_k_* and *Q_k_*, *_j_* are the distributions defined above.

Fig. 2c illustrates the computation of the KL divergence. The top panel shows the predictive distribution and normalized single-event likelihood as functions of *x_k_*. The middle panel shows their pointwise log-density ratio, log *P_k_* (*x_k_*)/*Q_k_*, *_j_* (*x_k_*). The bottom panel shows this ratio weighted by *P_k_* (*x_k_*), which, when integrated, gives the value of the KL divergence.

The KL divergence is the most familiar way to compare two distributions, and we include it as a natural point of reference. It is an asymmetric measure, meaning that, in general, *D*_KL_(*P* ‖ *Q*) ≠ *D*_KL_(*Q* ‖ *P*). We could also compute the KL divergence from the likelihood to the predictive distribution, *D*_KL_(*Q* ‖ *P*), or the average of these two divergences, 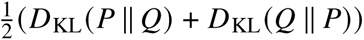, but both alternatives lead to a common issue, namely that the KL divergence can become arbitrarily large when its first argument places appreciable probability where its second argument assigns very little. Thus, *D*_KL_(*Q* ‖ *P*) can become large when an uninformative spike produces a broad likelihood. Choosing *D*_KL_(*P* ‖ *Q*) avoids that particular behavior but raises a complementary one: the divergence will grow large whenever the predictive distribution is broad relative to the likelihood, which can occur, for example, when informative spikes are sparse. We will see in the simulation and data analysis below that this behavior flags many spikes, even when their information is consistent with the accumulated information from past spiking.

## Simulation study

Validating local goodness-of-fit measures on real neural data is fundamentally limited: when the true state and the process that generated the spikes are unknown, we cannot tell whether flags reflect misfit, a latent state that departs from the animal’s physical position without any misfit, or the rare events that a correct model also produces, nor whether an absence of flags means that the model fits. We therefore first evaluate the three measures—HPD overlap, the rank-based predictive *p*-value, and KL divergence—in a simulation in which every departure from the decoder’s model is known by construction. We simulate a hippocampal place-cell population on a one-dimensional track and decode it with a Bayesian filter whose model matches the simulation. Three misfit windows break that match, each by perturbing either the observation model or the state-transition model. Two control windows present activity that departs from ordinary running—a replay sweep and sparse firing during immobility—but that the decoder’s observation model describes correctly. Because we know which part of the model is wrong in each window, we can determine how often each measure flags spikes under a correct model, which kinds of misfit it detects and which it misses, and whether it flags times when the latent state departs from the animal’s physical position without misfit, as in replay, or when the prediction is much broader than a spike’s likelihood, and interpret each result through our notion of consistency.

### Simulation design

#### Track and observation model

The state space is a one-dimensional linear track of length *L* = 100 arbitrary units (a.u.) discretized at unit spacing, *x* ∈ {0, 1, …, 100}. The simulated population consists of eleven place cells with Gaussian place fields centered at *μ_c_* ∈ {0, 10, …, 100} and shared standard deviation *σ*_pf_ = 10 a.u. The expected spike count of cell *c* per 1 ms time step at position *x* is

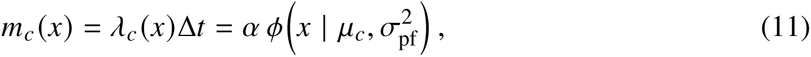

where Δ*t* = 1 ms, *φ*(· | *μ*, *σ*^2^) is the Gaussian density, and *α* = 5 is a scale parameter for the expected count. The peak expected count is 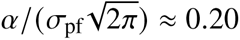 spikes per step, corresponding to a peak rate of approximately 200 Hz. Spike counts are drawn independently across cells at each step, *n_k_*_,*c*_ ∼ Poisson(*m_c_* (*x_k_*)). Except during the replay control described below, *x_k_* is both the animal’s physical position and the position represented by the simulated ensemble.

#### Position dynamics

Except where noted below, the animal’s position evolves as a Gaussian random walk with step standard deviation *σ*_pred_ = 0.5 a.u. The trajectory is reflected at 0 and 100 a.u. throughout the session, starts at *x*_0_ = 0, and is continuous across phases: the final position of each phase becomes the initial position of the next.

#### Decoder

The decoder is a Bayesian filter on the same grid of positions. Its model is the generative one everywhere except in the misfit windows and in the movement during the two controls. Its observation model treats the spike counts as independent Poisson variables with the expected counts of Eq. (11), using the true place fields; during each control, the expected counts are switched to the rates that generate that window’s spikes. Its state-transition model is a Gaussian random walk with the same step standard deviation *σ*_pred_, truncated at the track ends and renormalized. At each step, the decoder applies the transition to the previous filtering distribution to obtain the one-step predictive distribution *P_k_*, and then updates *P_k_* with the likelihood of all spike counts in the bin, including those of the cells that did not fire. The measures compare *P_k_* with the normalized single-event likelihood *Q_k_*, *_j_* of each spike in the bin.

#### Flag thresholds

A spike is flagged as a poor fit when its value on a measure crosses a threshold. The HPD-overlap and KL-divergence thresholds are set from the opening, well-specified part of the session (steps 0–6,000), with the values pooled across 100 independent realizations so that the thresholds are not dominated by the noise of a single run. Because low HPD overlap indicates disagreement, a spike is flagged when its value is at or below the 1st percentile of the pooled baseline values; because many baseline spikes are tied at that value, this rule flags 1.4% of baseline spikes rather than the nominal fraction. Because high KL divergence indicates disagreement, a spike is flagged when its value is at or above the 99th percentile. The rank-based predictive *p*-value instead uses a fixed threshold: a spike is flagged when *p* ≤ 0.05 (equivalently, − log(*p*) ≳ 3, with log the natural logarithm).

#### Phase schedule

The 32-second session (32,000 steps) contains three misfit windows and two controls, described below in the order in which they appear in Fig. 3a. Each misfit window perturbs the observation model *p*(*y_k_* | *x_k_*) or the state-transition model *p*(*x_k_* | *x_k_*_−1_), either in the generative process or in the decoder’s assumptions. The two controls change the spiking in ways the decoder’s observation model takes into account, so they contain no observation misfit; their movement—a deterministic sweep in one, an immobile animal in the other—is not a random walk. All other steps are well specified: the opening baseline, which sets the thresholds, and the three clean-recovery windows that follow the misfit windows (steps 10,000–14,000, 18,000–22,000, and 26,000–30,000), in which the generative process and the decoder return to the baseline configuration. Fig. 3b pools the recovery windows, without the replay steps, into a single well-specified column.

**Fig 3.**
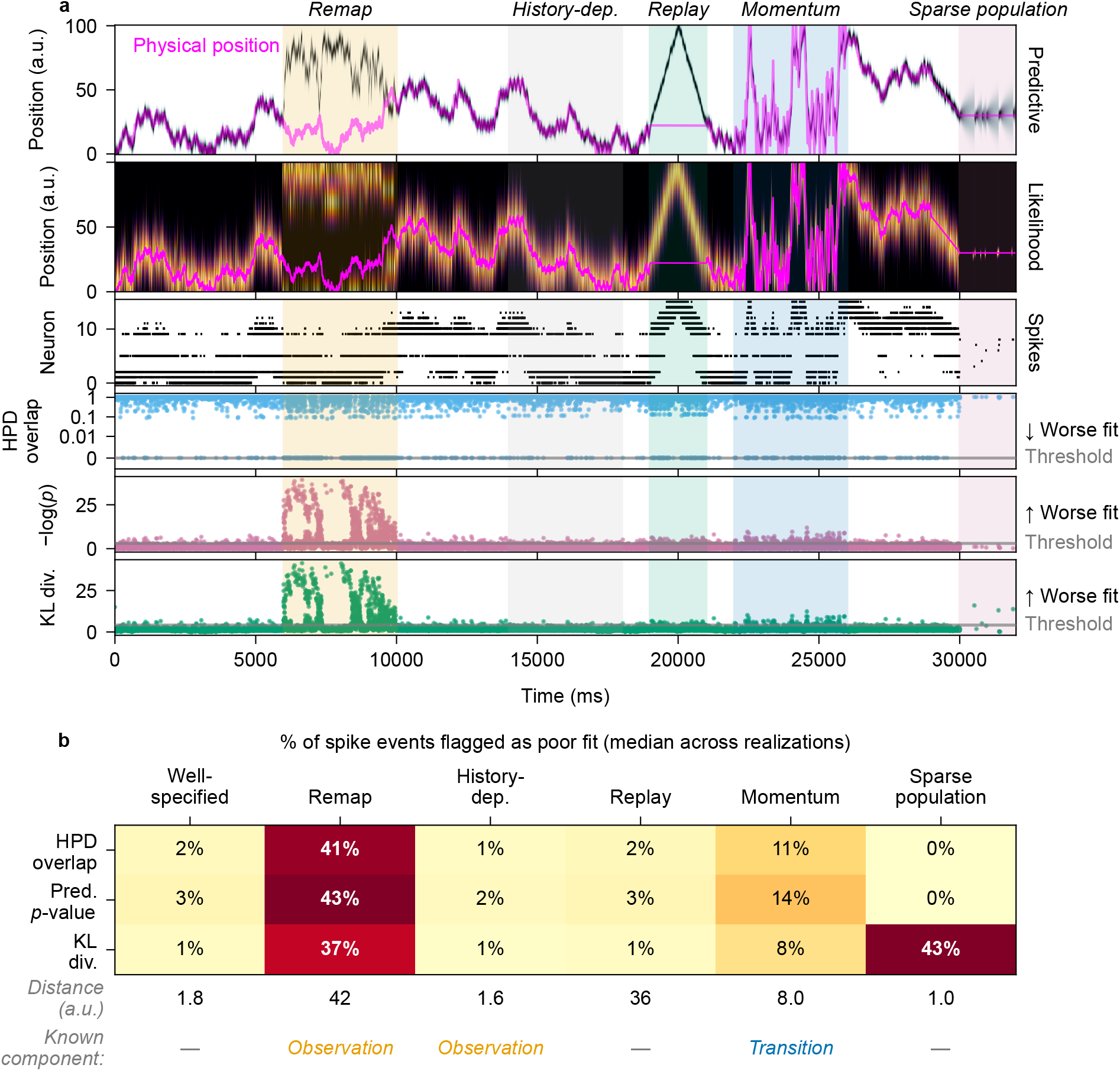
Per-spike goodness-of-fit measures for the simulation study. **(a)** Top row: the predictive distribution over time (1 ms per step) and position, with the animal’s physical position in magenta; shading marks the three misfit windows and two controls. Second row: the mean normalized per-spike likelihood in each spike-containing bin (averaged for display only). Third row: spikes of all sixteen neurons (eleven place cells, five sparse-population cells), sorted by field center. Bottom rows: HPD overlap (symmetric-log axis), −log(*p*) of the rank-based predictive *p*-value, and KL divergence for each spike event, with flag thresholds. **(b)** Median percentage of spike events flagged in each phase across 100 realizations, with the clean-recovery windows (excluding the replay event) pooled into the well-specified column. HPD and KL thresholds come from the pooled opening baseline (steps 0–6,000); the predictive *p*-value uses a fixed threshold of 0.05. The “Distance” row gives the median absolute distance (a.u.) between the decoded position (mean of the filtering distribution) and the animal’s physical position. The bottom row names the perturbed component (*Observation* or *Transition*); a dash marks well-specified and control phases.

#### Remap

During steps 6,000–10,000, the decoder-side observation model is perturbed. Spikes are generated from the true place fields, but the decoder evaluates them against a scrambled map: the eleven place-field centers are reassigned among the cells by a fixed random permutation that moves every field by at least three place-field spacings, modeling global remapping to statistically independent locations. The measures use the same scrambled map as the decoder and have no access to the true fields, so the remap is detectable only through its internal inconsistency: the scrambled map places spikes generated at neighboring positions at unrelated locations, which a smoothly moving state cannot explain.

#### History-dependent firing

During steps 14,000–18,000, the generative-side observation model is perturbed. Spike counts are generated with each cell’s Poisson mean set to zero for 1 ms after a step in which it spiked, followed by a 2 ms–10 ms burst window with a threefold rate increase, matching the rough phenomenology of CA1 pyramidal-cell refractory and burst windows; the decoder still assumes independent Poisson firing. Because the burst window outlasts the suppressed step, this modulation also raises each cell’s mean firing rate above the Poisson rate the decoder assumes. The per-spike spatial likelihood is unchanged, so the mismatch lies in the temporal joint distribution and overall rate of the spike train, not in the spatial information each spike carries.

#### Replay

During steps 19,000–21,000, within the second clean-recovery window, a replay event occurs. For this control, we distinguish the animal’s physical position, *z_k_*, from the internally represented position, *x_k_*, that drives the place-cell spiking. The animal is immobile, so *z_k_* remains fixed, while *x_k_* starts at *z_k_* and moves at constant speed to the farther end of the track and back. During the sweep the place cells fire at an elevated rate, with *α* = 20 in Eq. (11) (a peak rate of approximately 800 Hz). The decoder uses these elevated rates, so its observation model matches the replayed spikes. The replay steps are scored as their own column in Fig. 3b.

#### Momentum

During steps 22,000–26,000, the generative-side state-transition model is perturbed. The trajectory is generated with persistent velocity, *v_k_* = 0.88 *v_k_*_−1_ + *η_k_* and *x_k_* = *x_k_*_−1_ + *v_k_* with 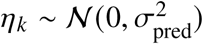; the decoder continues to assume a memoryless random walk with step standard deviation *σ*_pred_.

#### Sparse population

During steps 30,000–32,000, the animal is immobile at *x* = 30 a.u. It reaches this position at constant speed during the last second of the third clean-recovery window, so the control begins without an abrupt position jump; these approach steps remain in the well-specified column. The eleven place cells fall silent (their expected counts are set to zero), as can occur during immobility, and the only spikes come from a sparse population of five additional cells with narrow fields clustered near *x* = 30 a.u. (centers spread over 28.5–31.5 a.u., each with standard deviation 2 a.u.). Each fires as an independent Poisson process with a peak rate of 1 Hz, so the population produces only a few spikes per second and its spikes are separated in time. Before this window the same cells fire at a peak rate of 0.01 Hz. The decoder uses these phase-specific rates and knows that the place cells are silent, so its observation model is correctly specified.

### Simulation results

#### The measures rarely flag spikes when the model is correct

We first asked how often each measure flags spikes when the decoder’s model matches the process that generated the data. In the well-specified windows, HPD overlap flagged a median of 1.8% of spikes (interquartile range across realizations, IQR, 1.5–1.9%), the rank-based predictive *p*-value 2.8% (IQR 2.6–2.9%), and KL divergence 1.2% (IQR 1.0–1.3%), and the decoded position was a median of 1.8 a.u. (IQR 1.7–1.8 a.u.) from the animal’s position, less than one-fifth of the place-field standard deviation (Fig. 3b). These windows were not used to set the thresholds, so this rate is what we would expect from the measures on well-fit data.

#### All three measures detect an incorrect place-field map

We next asked whether the measures could detect an incorrect observation model when they, like the decoder, have access only to that model. When the decoder used the scrambled map, successive spikes, generated from neighboring positions, had likelihoods centered on unrelated locations, and each spike’s likelihood often fell outside the region supported by the prediction (Fig. 3a). HPD overlap flagged 41% of spikes (IQR 22–57%), the predictive *p*-value 43% (IQR 23–57%), and KL divergence 37% (IQR 19–52%), and the decoding error rose to 42 a.u. (IQR 25–65 a.u.; Fig. 3b). These percentages varied widely across realizations because they depend on how much of the scrambled map the trajectory visits. The measures thus identified the period in which the observation model was wrong and the decoded position unreliable.

#### Misfit that leaves spatial information intact is not flagged

Not every departure from the model produces conflicting information about position. When cells fired with refractory periods and bursts that the decoder did not model, the spatial tuning of each spike was unchanged, so each spike’s likelihood was still centered near the animal’s location. The decoder treats the spikes within a burst as independent evidence, so its predictive distribution narrowed, but it did not move away from the true position. HPD overlap flagged only 1.0% of spikes (IQR 0.7–1.3%), the predictive *p*-value 1.5% (IQR 1.3–1.7%), and KL divergence 0.6% (IQR 0.4–0.7%), at or below the well-specified rates, and the decoding error was essentially unchanged (1.6 a.u.; IQR 1.5–1.7 a.u.; Fig. 3b). The measures thus did not flag every form of misspecification, but their silence here coincided with an accurate decoded position.

#### Replay is not mistaken for model failure

In the hippocampus, the decoded position departs from the animal’s location during replay [8], and a useful measure should not treat these events as errors. Because the simulated sweep began at the animal’s position and moved by only a fraction of the random walk’s step standard deviation per step, the prediction could follow it continuously far from the animal, and each spike remained consistent with it (Fig. 3a). HPD overlap flagged 2.2% of spikes (IQR 1.9–2.3%), the predictive *p*-value 2.8% (IQR 2.6–3.0%), and KL divergence 1.3% (IQR 1.2–1.4%), similar to the well-specified windows, even though the decoded position was a median of 36 a.u. (IQR 32–40 a.u.) from the animal (Fig. 3b). Because the measures assess whether spikes are consistent with the model’s prediction rather than with the animal’s physical position, they leave replay unflagged.

#### The measures also detect an incorrect movement model

When the animal moved with momentum that the decoder’s random walk did not capture, the prediction lagged behind the animal. HPD overlap flagged 11% of spikes (IQR 10–11%), the predictive *p*-value 14% (IQR 13–15%), and KL divergence 8.5% (IQR 7.8–9.3%): several times each measure’s well-specified rate, though less than in the remap window. The median decoding error rose more than fourfold, to 8.0 a.u. (IQR 7.5–8.4 a.u.; Fig. 3b). The flag rate is modest relative to this error because a single spike localizes the animal only coarsely: its likelihood extends well beyond the lag, so most spikes still overlap the offset prediction, and only spikes from cells whose fields lie far from the predicted position are flagged.

#### Only KL divergence flags consistent sparse firing

Finally, we asked whether the measures distinguish between two cases: a likelihood that is narrower than the prediction but lies within it, and a likelihood that lies outside it. When the animal was immobile and only the sparse population fired, the prediction broadened between isolated spikes, and each spike’s narrow likelihood fell within the range the prediction supported (Fig. 3a). HPD overlap flagged 0.0% of spikes (IQR 0.0–0.0%) and the predictive *p*-value 0.0% (IQR 0.0–0.0%), whereas KL divergence flagged 43% (IQR 36–57%), even though decoding remained accurate (median error 1.0 a.u.; IQR 0.78–1.5 a.u.; Fig. 3b). KL divergence responds to the difference in concentration between the two distributions, not to a conflict in the information they carry about position.

Together, these results show that HPD overlap and the predictive *p*-value detect misfit that places a spike’s likelihood outside the region supported by the prediction: they flagged many spikes during remapping and fewer during momentum. They did not flag history-dependent firing, which left each spike’s likelihood centered near the animal, and they did not mistake replay for model failure, even though the decoded position was far from the animal. KL divergence differed from them only in the sparse-population control, where it flagged spikes whose likelihood was narrower than the prediction but lay within it, while HPD overlap and the *p*-value flagged almost none.

## Real hippocampal data analysis

### Data collection and behavioral task

We applied the local goodness-of-fit measures to units recorded from the rat hippocampus during one 24-minute recording epoch, in which the animal performed a spatial bandit task on a maze with three radially arranged Y-shaped foraging patches extending from a central point (Fig. 4, inset). The details of this task and the data collection methods are described by Comrie et al. [18]. Briefly, the rat was trained to run between six reward ports located in three Y-shaped “patches”. Each port was assigned a reward probability of 0.2, 0.5, or 0.8, such that one patch was best on average. Reward probabilities changed periodically after a preset time or trial limit. Recordings were made with 22 tetrodes advanced to CA1; two additional tetrodes, one in each hemisphere, served as local references in the corpus callosum.

**Fig 4.**
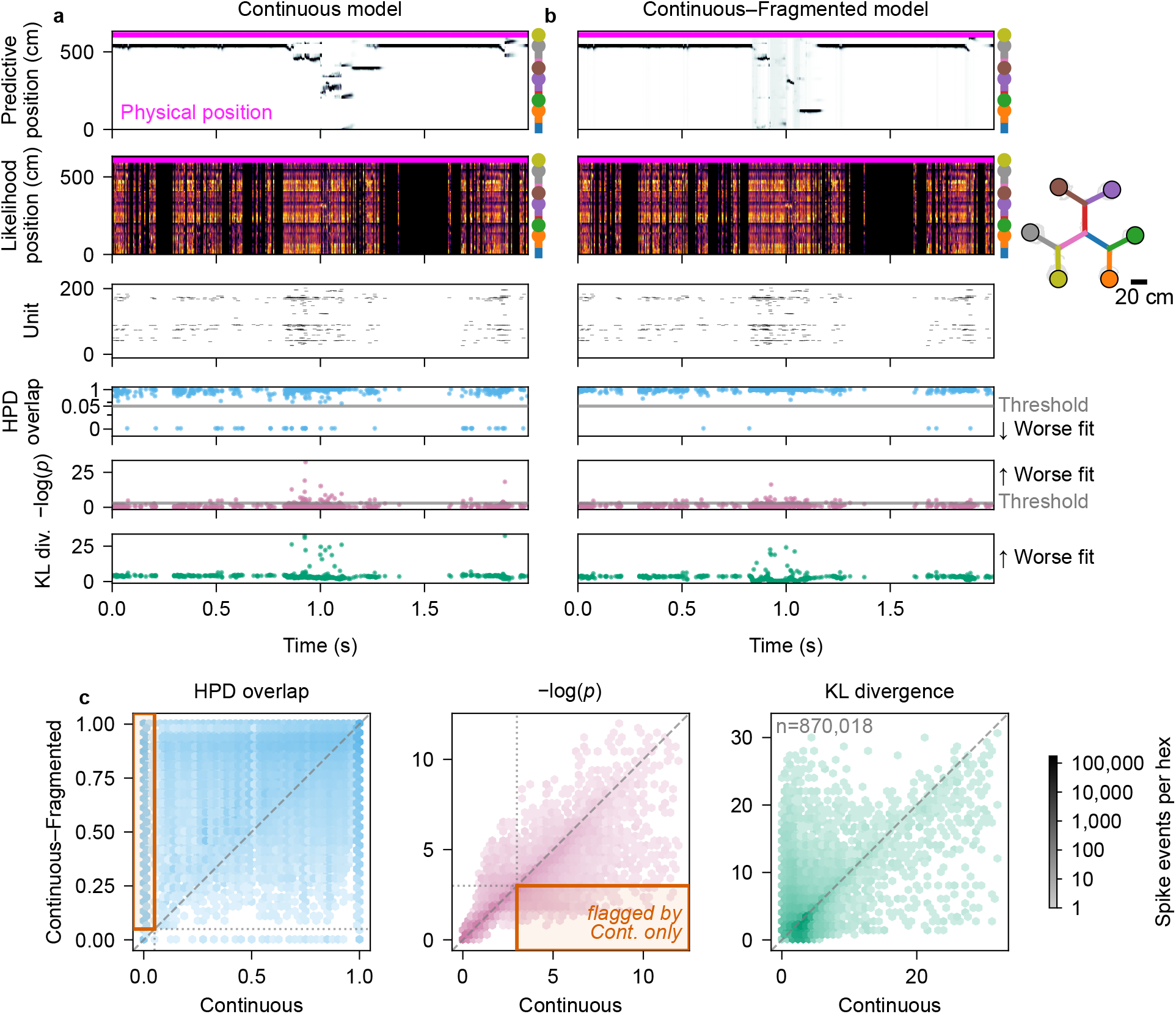
Per-spike goodness-of-fit measures applied to a hippocampal recording during navigation on a multi-arm maze (203 sorted units, 182 of which fired in this epoch; data from Comrie et al. [18]), evaluated under a Continuous and a Continuous–Fragmented state-space model. The inset shows the two-dimensional maze with the animal’s trajectory in gray; colored dots mark the six reward ports, and segment colors match the bars beside rows 1–2. **(a, b)** A window containing a candidate replay event, decoded with the Continuous (a) and Continuous–Fragmented (b) models; time is relative to the start of the window. Row 1: the one-step predictive distribution over linearized position, with the animal’s physical position overlaid. Row 2: the mean normalized per-spike likelihood in each spike-containing bin (averaged for display only), identical under both models. Row 3: the spike raster, with units sorted by place-field peak position. Rows 4–6: HPD overlap (symmetric-log axis), −log(*p*), and KL divergence for each spike event. Horizontal lines mark the flag thresholds, 0.05 for HPD overlap and −log(0.05) 3 for the predictive *p*-value; KL divergence has none. **(c)**Per-spike values across the epoch under the Continuous (horizontal) and Continuous–Fragmented (vertical) models; the dashed line is the identity. Dotted lines mark flag thresholds; outlined boxes contain spikes flagged only under the Continuous model. Color gives spike events per hexagon (shared log scale); *n* is their total; the − log(*p*) axis omits a few larger values.

For the present analysis, we sorted the continuous 30 kHz neural signal into putative single units with MountainSort4 [19]. In the MountainSort4 pipeline, the signal was band-pass filtered and whitened during preprocessing; negative-going events were then detected on each tetrode at a threshold of 3 in whitened units, with a minimum inter-event interval of 10 samples (≈0.33 ms). Units were retained from automated curation only, without manual cluster curation. Because curation was automated, some units may contain multiunit activity or represent incompletely merged clusters; this does not affect our purpose here, which is to demonstrate the measures rather than to characterize individual neurons.

For decoding, we linearized the animal’s position by projecting it onto the nearest point of a graph of the track’s segments, discretized linear position into bins of approximately 2 cm, and binned spikes at 2 ms resolution. Both decoders used the same observation model. Each unit’s place field was estimated by Gaussian kernel density estimation as the unit’s mean firing rate multiplied by the ratio of the smoothed density of the animal’s positions at the unit’s spikes to the smoothed occupancy density, with a kernel standard deviation of with 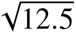 ≈ 3.5 cm. Both decoders started from a uniform distribution over positions on the track.

The two decoders differ only in how they assume the represented position changes over time. The Continuous model assumes that it changes gradually: from one time bin to the next it takes a Gaussian random-walk step along the track with variance 6.0 cm^2^. The Continuous–Fragmented model adds a second, Fragmented mode, in which the represented position in each time bin is drawn uniformly from the whole track, independent of its previous value, so that it can jump to a distant location in a single step [2]. The model starts in the Continuous or Fragmented mode with probability 0.5 or 0.5, and in each time bin it stays in the Continuous mode with probability 0.98 (switching to Fragmented with probability 0.02) and in the Fragmented mode with probability 0.98 (switching back with probability 0.02). Within the Continuous mode, position follows the same random walk as in the Continuous model, except in the first time bin after the model leaves the Fragmented mode, when the position is again drawn uniformly.

The Continuous–Fragmented decoder represented the joint state of discrete dynamics mode and position, with the same position-dependent observation model in both modes. For all three measures, we marginalized its one-step predictive distribution over dynamics mode and compared the resulting distribution over position with one copy of the shared observation likelihood. The measures for both decoders were therefore defined on the same position grid. The rank-based predictive *p*-value used the same fixed threshold as in the simulation, flagging *p* ≤ 0.05. For HPD overlap, the recording has no designated clean baseline period from which to derive a percentile threshold, so we flagged values at or below a fixed threshold of 0.05. We did not threshold KL divergence, whose simulation threshold also came from a baseline period; we instead compare its values between the two models.

### Data analysis results

#### The Continuous model is flagged more often during immobility

A standard decoder assumes that the hippocampal representation moves continuously. During immobility, however, a replayed trajectory need not begin at the animal’s location: the representation can jump to a distant part of the environment and replay a path from there [20]. Unlike the simulated replay, which began at the animal and moved slowly enough for a random walk to follow, such an event places spikes’ likelihoods far from a prediction that can move only gradually. We therefore expected the Continuous model to be locally inconsistent with the spiking during these events. We compared it with the Continuous–Fragmented model, whose Fragmented mode lets the representation jump to any location; we previously used a related model to capture replay and other nonlocal hippocampal representations [2]. The two models share one observation model, so each spike’s likelihood is identical under both, and any difference in the measures comes from the prediction. Consistent with this expectation, the Continuous model’s flag rates were higher while the animal was immobile (speed below 4 cm/s) than while it moved: 4.4% against 1.2% for HPD overlap and 4.1% against 3.8% for the predictive *p*-value. KL divergence was also higher during immobility (median 3.5 against 2.9), but this difference need not reflect misfit: in the simulation’s sparse-population control, KL divergence flagged many spikes during immobility because the prediction broadened between isolated spikes, even though each spike was consistent with it.

#### Allowing jumps removes most HPD-overlap flags in a candidate replay event

In a 2 s window containing a candidate replay event, the animal was at rest (median speed 0.86 cm/s) and successive spikes had likelihoods centered on distant maze locations (Fig. 4a). The Continuous model’s prediction could move only gradually and did not keep up with them, so for some spikes it concentrated where the spike gave no support: of the window’s 649 spikes, HPD overlap flagged 43 and the predictive *p*-value 45. The Continuous–Fragmented model reserves part of its prediction for a jump to any location. During this event, its predicted probability of the Fragmented mode rose from a median of 0.047 to a peak of 0.96, exceeding 0.8 for a total of 54 ms (Fig. 4b). Under this model, HPD overlap flagged 5 spikes and the predictive *p*-value 33, and 448 of the 649 spikes had larger *p*-values than under the Continuous model. A broad prediction does not make a spike expected; it only makes the spike less surprising than under a prediction concentrated at the wrong location. KL divergence also fell for most spikes in the window (median 3.2 to 2.2).

#### Across the epoch, allowing jumps removes most HPD-overlap flags

The same pattern held across all 870,018 spike events in the 24-minute epoch (Fig. 4c). HPD overlap flagged 2.2% of spikes under the Continuous model and 0.2% under the Continuous–Fragmented model. All but 61 of the 18,790 spikes flagged under the Continuous model had an overlap of zero: the spike’s likelihood and the prediction had disjoint HPD regions. Of the spikes flagged under the Continuous model, 17,289 (92%) were not flagged under the Continuous–Fragmented model (*rescued*), whereas only 176 spikes were flagged under the Continuous–Fragmented model alone (*newly flagged*).

#### The predictive *p*-value points the same way, more weakly

The predictive *p*-value rescued fewer spikes, and its flag rate fell only from 3.9% to 3.0%. Of the 33,954 spikes it flagged under the Continuous model, 9,373 (28%) were rescued and 1,706 were newly flagged.

#### KL divergence provides only weak evidence

KL divergence fell under the Continuous– Fragmented model for 94% of spikes (median 3.1 to 2.2), the same direction as the other two measures. For the remaining 6%, which started low (median 1.5), KL divergence rose, usually modestly (median 2.2), although a few values reached 30. This rise matches the simulation’s sparse-population control, in which KL divergence flagged spikes whose narrow likelihood lay within a broader prediction: the Fragmented mode broadens the prediction, which can raise KL divergence for a spike that was already consistent with it. Because KL divergence responds to how broad the prediction is as well as to where it is concentrated, its changes here are weak evidence about fit.

Together, these results show that the measures can guide a modeling decision on real data. The Continuous model’s HPD-overlap flags were concentrated where a random walk is expected to fail, during immobility, when the hippocampal representation can jump, and adding a mode that allows such jumps removed most of them while creating few new ones. The predictive *p*-value agreed, more weakly. KL divergence changed in the same direction for most spikes, but because it also responds to the breadth of the prediction, it adds little independent evidence.

## Discussion

State-space modeling has become a powerful tool for describing the latent dynamics of neural systems. Validating these models is inherently difficult because latent states are unobservable and many models are purposefully simplified. To address this, we developed local goodness-of-fit measures that compare the information carried by each spike with the information the model carries forward from all preceding spikes. Rather than accepting or rejecting a model in its entirety, these measures locate the times when the two are inconsistent. We found that two of them, HPD overlap and the rank-based predictive *p*-value, flag spikes whose likelihood lies outside the region supported by the prediction but not spikes from times when the latent state departs from the animal’s physical position without misfit, as in replay, and that in a hippocampal recording they point, HPD overlap most clearly, to a specific refinement of the decoding model.

HPD overlap and the rank-based predictive *p*-value behaved as a consistency measure should. In the simulation, they flagged spikes whose likelihood lay outside the region supported by the prediction, as when the place-field map was wrong. They did not flag spikes whose likelihood lay within a broad prediction, and they were not triggered when the decoded position departed from the animal during replay. HPD overlap provides a geometric measure of consistency that remains interpretable when one high-density region contains another; the rank-based predictive *p*-value provides a direct statistical assessment of each observation. The two can nonetheless differ. In the recorded data, the predictive *p*-value’s flags were less concentrated in periods of immobility than HPD overlap’s and changed less when the model was refined, so the two measures are best read together.

KL divergence is computationally simple but sensitive to the tails of the distributions, and it has no clear upper bound, which makes a threshold difficult to set. In the simulation, it also flagged spikes whose narrow likelihood lay within a much broader prediction, which the other two measures treated as consistent. KL divergence is one member of the larger family of *f* -divergences, which also includes the total variation distance and the Pearson *χ*^2^ divergence [21]. Substituting another member would emphasize different features of the prediction–likelihood mismatch, but because these measures all target distributional difference rather than consistency, this behavior would likely carry over to the family as a whole. This behavior is a liability for consistency assessment, but it could be useful where the goal is to flag times when the prediction is uncertain relative to the information carried by individual spikes. When the goal is consistency in estimates of the state process, we recommend HPD overlap and the rank-based predictive *p*-value as primary diagnostic measures.

In practice, HPD overlap and the predictive *p*-value make it possible to identify specific periods when a state-space model is inconsistent with the spiking data, and to turn that diagnosis into a concrete modeling decision. Consider a researcher decoding the hippocampal recording with the Continuous model alone. Its HPD-overlap flags are concentrated in periods of immobility, when the hippocampal representation can jump to distant locations, and during the candidate replay event its prediction cannot keep up with spikes whose likelihoods move between distant parts of the maze. This pattern suggests that the continuous transition model is too restrictive. Consistent with that hypothesis, adding a Fragmented mode that permits discrete jumps eliminates the HPD-overlap flag for 92% of spikes flagged under the Continuous model across the recording epoch while creating few new flags (Fig. 4c). A single score for the whole epoch could rank the two models, but it would not identify which individual observations drive the difference. Because a broader prediction is also more permissive, a fall in flags under an alternative model should be read alongside how broad its prediction is. The Continuous–Fragmented model, for example, reserves some probability for a jump to any location at every step. Here, the local measures, alternative-model comparison, and scientific context together support a specific refinement of the model. More broadly, this diagnose-then-refine cycle enables a feedback loop between model development and assessment that can improve the quality of state-space models.

The simulation also makes two limits of these measures concrete. First, each spike is compared with the prediction through its own normalized likelihood, which excludes silent neurons and the total spike count (see Local goodness-of-fit measures), so the measures cannot see misfit that leaves each spike’s likelihood in place. History-dependent firing in the simulation changed the timing and rate of spikes but not the spatial information each carried, and it went unflagged; misfit of this kind requires a complementary count or time-rescaling test [6, 22]. Second, the measures operate at the time scale of single spikes. Individual place-cell spikes are informative enough that we often detected local misfit at this scale, but a single spike localizes the animal only coarsely, so a persistent lag between the prediction and the animal, as in the simulated momentum window, was flagged in only a modest fraction of spikes. Such misfit, and local errors in neural systems whose individual observations are less informative, may be easier to detect by aggregating the measures over longer intervals; choosing those intervals may require prior knowledge about the system or exploring the measures across several time scales.

We illustrated these measures on simulated and real spike-sorted data from rat hippocampus, but they extend to other neural data types modeled with state-space models. In particular, clusterless decoding has been shown to give lower decoding error and better-calibrated uncertainty than decoding from spike-sorted units [5]. As defined in the Methods section, the measures apply directly to clusterless models. In that case, a flagged spike indicates that a spike with that waveform was unexpected given the information the model had accumulated up to that time.

A second extension concerns the distribution each spike is compared with. For simplicity, we compared each spike’s likelihood with the one-step predictive density, which aggregates information from the past. The predictive density could be replaced by any posterior density from the state-space model. A smoothing density (e.g., from a fixed-interval smoother over the observation period), for example, would use both past and future data and could increase the power to detect model misspecification. Because filtering and smoothing densities also condition on the spike being assessed, that spike’s own contribution would first need to be removed, for example by leave-one-out smoothing, so that the spike is not compared against a distribution it helped form.

Applying these measures more broadly also raises computational challenges. The cost of these measures increases as the dimensionality of the state process increases. In all of our examples, we computed KL divergence and HPD overlap by partitioning the state space into a lattice and computing the likelihoods and predictive densities at each point in that lattice. As the dimension of the state increases, the size of that lattice grows exponentially. We also evaluated the predictive checks exactly: for spike-sorted data on a discretized state space, the *p*-value is a finite sum over the recorded neurons and is inexpensive whatever the state-transition model. For continuous marks, as in clusterless decoding, the predictive check must instead be approximated by sampling from the predictive state and observation distributions. For observation spaces that are finite but large, sampling could also avoid enumerating every outcome; whether this is more efficient will depend on the model and the required Monte Carlo precision. Sampling-based estimators could likewise improve the tractability of KL divergence and HPD overlap in moderate dimensions, although the number of samples required for accurate estimates will still grow with dimension. Alternatively, the measures could be computed in lower-dimensional subspaces of the state space; identifying subspaces that retain the statistical power of the full space remains for future work.

To make these measures straightforward to apply in other studies, we provide them as an open-source Python package, `statespacecheck` (https://github.com/edeno/statespacecheck). For a spike-sorted decoder, the package needs only quantities that a grid-based decoder already computes: the one-step predictive distribution in each time bin, each neuron’s firing rate as a function of the state, and the time bin and neuron of each spike. From these it returns the HPD overlap, the rank-based predictive *p*-value (evaluated exactly over the recorded neurons), and the KL divergence for every spike, along with the building blocks (the normalized single-event likelihood and the predictive distribution over neurons) and a helper for setting thresholds from a baseline period. For clusterless decoders, the package computes the same three measures from the joint mark intensity *λ*(*y*, *x_k_*), approximating the predictive *p*-value by Monte Carlo as in Fig. 2b. The package does not depend on how the decoder was fit, so it can be applied to an existing model without modification. Because the measures yield one value per spike, they are most informative alongside the decoded state. The analysis code includes a desktop viewer that shows the per-spike measures over a full decoded recording epoch and lets the user compare any spike’s likelihood with the prediction at that time. A companion website (https://edeno.github.io/statespacecheck-paper/) shows how the measures are computed and lets readers inspect the simulated and hippocampal examples.

We believe that these local goodness-of-fit tools provide an important step toward making inferences from state-space neural models more reliable and interpretable. By pinpointing specific times when the model assumptions break down, researchers can be more confident in their inferences. Ultimately, these measures empower us to refine our models of neural dynamics and deepen our understanding of how the brain encodes and processes information.

## Ethical statement

The recording analyzed here was collected for a previous study [18]. All animal procedures were approved by the Institutional Animal Care and Use Committee at the University of California, San Francisco (protocol AN174991).

## Acknowledgements

This work was supported by the National Institutes of Health (RF1MH130623 to U.T.E. and L.M.F. and R01MH136875 to L.M.F.), the Simons Collaboration on the Global Brain (542971 to U.T.E. and 521921 and 542981 to L.M.F.), and the Howard Hughes Medical Institute (L.M.F.). A.E.C. was supported by the National Science Foundation Graduate Research Fellowship Program (1650113), the National Institutes of Health (F31MH124366), and the UCSF Jonas Cohler Discovery Fellowship. The authors used Claude (Anthropic; Claude Opus 4.8, 5.0, and 5.5) to assist with writing and reviewing the simulation, analysis, and figure-generation code and with editing the manuscript text.

The authors reviewed and revised all AI-assisted material and take full responsibility for the content of this article.

The authors declare that they have no competing interests.

## Data availability statement

The hippocampal recording analyzed here was originally reported by Comrie et al. [18] and is available on the Distributed Archives for Neurophysiology Data Integration (DANDI) Archive as dandiset 001942 (https://dandiarchive.org/dandiset/001942). The decoder inputs for the recording epoch analyzed in Fig. 4—the animal’s position, the spike times of the sorted hippocampal units, and the track geometry—are archived on Zenodo (doi:10.5281/zenodo.23020757).

## Code availability

The local goodness-of-fit measures are implemented in the statespacecheck Python pack-age, available at https://github.com/edeno/statespacecheck and installable from the Python Package Index (pip install statespacecheck); the analyses in this paper used version 0.3.1, archived on Zenodo (doi:10.5281/zenodo.23011969). All simulation, analysis, and figure-generation code, together with the interactive viewer and the source of the project website (https://edeno.github.io/statespacecheck-paper/), is publicly available at https://github.com/edeno/statespacecheck-paper and archived on Zenodo (doi:10.5281/zenodo.24116403).

## Author contributions

**Conceptualization:** UTE, SZ, ELD

**Methodology:** UTE, SZ, ELD

**Software:** SZ, ELD.

**Validation:** ELD.

**Formal analysis:** SZ, ELD

**Investigation:** SZ, UTE, ELD

**Data curation:** AEC, ELD.

**Resources:** ELD.

**Writing – original draft:** SZ, UTE, ELD.

**Writing – review & editing:** SZ, AEC, LMF, UTE, ELD.

**Visualization:** SZ, ELD.

**Supervision:** UTE.

**Funding acquisition:** UTE, LMF.

